# Neuronal dysfunctions and cognitive deficits in a multi-hit rat model following cumulative effects of early life stressors

**DOI:** 10.1101/2020.06.09.141754

**Authors:** Tiyasha Sarkar, Nisha Patro, Ishan Kumar Patro

## Abstract

Perinatal protein malnourishment is a leading cause for mental and physical retardation in children with poor socioeconomic conditions. Such malnourished children are vulnerable to additional stressors, that may synergistically act to cause neurological disorders at adulthood. In this study, the above mentioned condition is mimicked via a multi-hit rat model in which pups born to protein malnourished mothers (LP) were co-injected with polyinosinic:polycytidylic acid (Poly I:C; viral mimetic) at Postnatal day (PND) 3 and lipopolysaccharide (LPS; bacterial mimetic) at PND 9. Individual exposure of Poly I:C and LPS was also given to LP pups to correlate chronicity of stress. Similar treatments were also given to control pups. Hippocampal cellular apoptosis, β III tubulin catastrophe, altered neuronal profiling and spatial memory impairments were assessed at PND 180, using specific immunohistochemical markers (active caspase 3, β III tubulin, doublecortin), Golgi studies and cognitive mazes (Morris Water Maze and T maze). Increase in cellular apoptosis, loss of dendritic arborization and spatial memory impairments were higher in multi-hit group, than the single-hit groups. Such impairments observed due to multi-hit stress, mimic conditions similar to many neurological disorders and hence it is hypothesized that later life neurological disorders might be an outcome of multiple early life hits.

**Summary Statement:** This study is first of its kind which practically studies the combined effects of major early life stressors like protein malnourishment, viral and bacterial infections on the nervous system.

## Introduction

Approximately, 4 million neonatal deaths occur every year due to social crisis and 99% of such neonates belong to underdeveloped countries (Malqvist 2011). A child that survives poverty and social crisis might be at a higher risk for developing later life neurological disorders and hence, the concept of perinatal multi-stress emerges according to which, the children that encounter stress during perinatal period in the form of malnourishment, viral and bacterial infection, parental separation and abuse are vulnerable to development of neuropsychiatric disorders (Lai and Huang, 2011; Hoeijmakers et al., 2015; Syed and Nemeroff, 2017; Malinovskaya et al., 2018; Sarkar et al., 2019). The UNICEF data 2018, projects that 50% of death among neonates is due to undernutrition and additionally, nutritional inadequacy is also attributable to severity and death caused by common infections by delaying the recovery process of body. Again, according to world hunger index among different type of malnourishment, protein malnourishment is majorly responsible for retardation, delayed physical and mental growth and child death. Furthermore, viral and bacterial infectionsare another common issue in already malnourished children belonging to the poorer community because of the unhygienic surroundings in which the children grow (Rytter et al., 2014; Walson and Berkley, 2018). All these stressed conditions can together modulate the CNS and may be responsible for causing later life brain adversities.

Protein malnourishment, Poly I:C (double stranded RNA used to create a viral mimic animal model) and LPS, (bacterial endotoxin, widely used as a bacterial infection agent) are reported to change the neuronal architecture of fetal brain by destabilizing the synaptic connectivity leading to weakening the process of memory formation, retention and consolidation capacity in rats (Nishi et al., 2010; Yirmiya et al., 2011; Alamy and Bengelloun, 2012; Naik et al., 2015; Sinha et al., 2018). Both malnourishment and immune inflammation during early age play deleterious role via interacting with cellular integrity of the brain, hence multi stress together might increase the chances of brain deterioration by many folds. Connectivity failure in neuronal circuitry is one of the consequences of early life stress exposure, caused due to deformities of residential hippocampal neurons that further lead to memory wreckage in stressed individuals (Aas et al., 2012) which is very much common in case of neurological disorders like Alzheimer’s, Schizophrenia, autism etc. (Taylor et al., 2014; Dalle and Mabandla, 2018; Justice, 2018).

Decline in the number of viable neurons by caspase mediated apoptotic pathways are common in stress induced models (Moleur et al., 1998; Watanabe et al., 2002). Caspases are main executer cysteine proteases, that when activated, nucleophilically attack targeted proteins resulting in cleavage and cellular apoptosis. Neuronal degeneration could be due to stress mediated apoptotic pathways during which activated caspases cleaves cytoskeletal and integral cellular proteins, fragmenting and decreasing the overall cell number in brain. Cytoskeletal proteins are the building blocks of cellular extensions including neuronal arborization and are crucial for stable synapse formation, proper translation, transport and alignment of cytoskeletal subunits (Koleske, 2013). β III tubulin is a subclass of tubulin family encoded by TUBB3 gene in humans, specific to neurons and can be used as a marker for mature neuronal profiling. Neuronal damage can be identified as fragmentation and disruption of β III tubulin subunits leading to loss of dendritic arborization and connectivity (Poirier et al., 2010; Verstraelen et al., 2017), further heading an affected individual towards neurological disorders like Alzheimer’s, Autism, ADHD and Schizophrenia (Bishop et al., 2010). Among neuronal sub types, pyramidal neurons are associated with memory functioning (Kasai et al., 2010) and during memory impairment pyramidal neurons of CA1 (CornuAmmonis) and CA3 regions of hippocampus are largely found to be affected (McEwen et al., 2016). The foremost effects of stress on neurons was identified as alterations in the classical morphology of the neurons that appeared either stubbed, confined or extra-long types. In the stress induced brains, there is also an overall increase in dendritic fragmentations and neuronal death. Decrease in the arborization of neurons due to cell death and fragmentation causes connectivity failure interrupting synapse formation which is a common cause for memory impairment and neurological abnormalities (Kulkarni and Firestein, 2012).

Loss of mature neurons also calls for formation of new neurons (Kuhn et al., 2015; Quadrato et al., 2014). New neurons during adult neurogenesis are formed in the Dentate Gyrus (DG) in all mammalian species including humans (Synder et al., 2011). The newly formed immature neurons subsequently migrate and integrate into the circuitry on demand i.e., when and where necessary. Naive neurons are initially high in excitability and low in inhibition but, after maturation and integration in the functional layers, they become stable and start extending their processes, forming new arborizations and connections with the residential local neurons (Hastings and Gould, 1999; Dieni et al., 2016). Doublecortin (DCX), a neuronal migratory microtubule protein encoded by DCX gene is specific for naive and migratory neurons (Brown et al., 2003). Stress induced changes in adult neurogenesis have been reported by many researchers but relation between neuronal disintegration, adult neurogenesis, neuronal migration and memory impairment has not been well established.

The overall cellular malfunctioning in hippocampus due to various early life stressors may finally lead to memory impairments, which can be considered as onset of neuropsychiatric disorders. Hippocampus is associated with spatial memory impairments (Broadbent et al., 2004; Shrager et al., 2007) which includes both spatial reference and spatial working memory (Niewoehner et al., 2007; Bizon et al., 2012). Spatial reference memory can be differentiated from spatial working memory as the former allows left right discrimination whereas the latter is used to keep an account of recent activities like completing a task or understanding a sentence (Cowan, 2014). Hippocampus was initially found to be responsible only for declarative or long-term memory but recent studies have supported the involvement of hippocampus with working memory retention (Jeneson et al., 2011; Leszczynski, 2011; Yee et al., 2014). Such is the need for cognition in normal living conditions and the impairment of cognitive abilities can directly be associated with neuropsychiatric disorders (Lajud and Torner, 2015).

Most stress-oriented studies deal with a single type of stressor. Early developmental windows are prone to every minute changes occurring in the environment and hence the developing period is susceptible to multiple stressors which cumulatively may act on the developing brain causing later life abnormalities. Moreover, the synergistic and coordinated action of multiple early life stressors on the developing brain is highly challenging and require utmost attention. In this study, maternal malnourishment is used to create a stressed uterine environment and the F_1_ pups born from both healthy and malnourished mothers were further subjected to viral and bacterial infections with either polyinosinic:polycytidylic acid (Poly I:C) or Lipopolysaccharide (LPS) or both simultaneously. As malnourished subjects might be susceptible to immune infection, such multi-hit models depict the scenario relatable to most of the underdeveloped countries. Altered morphology and degeneration of neurons, in the rat hippocampus, was studied in context to memory impairment, using specific immunohistochemical markers and memory oriented cognitive mazes.

## Results

### Early life stressors (LP, Poly I:C and LPS) singularly or in combination induced cell death and cell layer damage in adult rats (PND180), showing cellular changes that are prominent and common in neurodegenerative disorders

Active caspase 3 positive intensely labeled cells were seen in the representative immunohistochemical images of different treated groups (red arrows). In perinatally LP fed animals there was an upregulation of active caspase 3 (Fig. 2E) when compared to HP control (Fig. 2A) and when LP animals were administered with single dose of either Poly I:C or LPS (LP+Poly I:C and LP+LPS), the active caspase 3 expression was further exaggerated (Figs. 2F,G respectively), increasing the number of cells undergoing apoptosis. Furthermore, when both Poly I:C and LPS were administered simultaneously to LP animals (LP+Poly I:C+LPS group), a vigorous increase in active caspase 3 positivity was recorded (Fig. 2H). Similar treatments were also administered to the HP animals (Figs. 2B, C, D) but the extent of cellular damage in HPgroupswere much less as compared totheir respective LP groups.

**Fig. 1.**
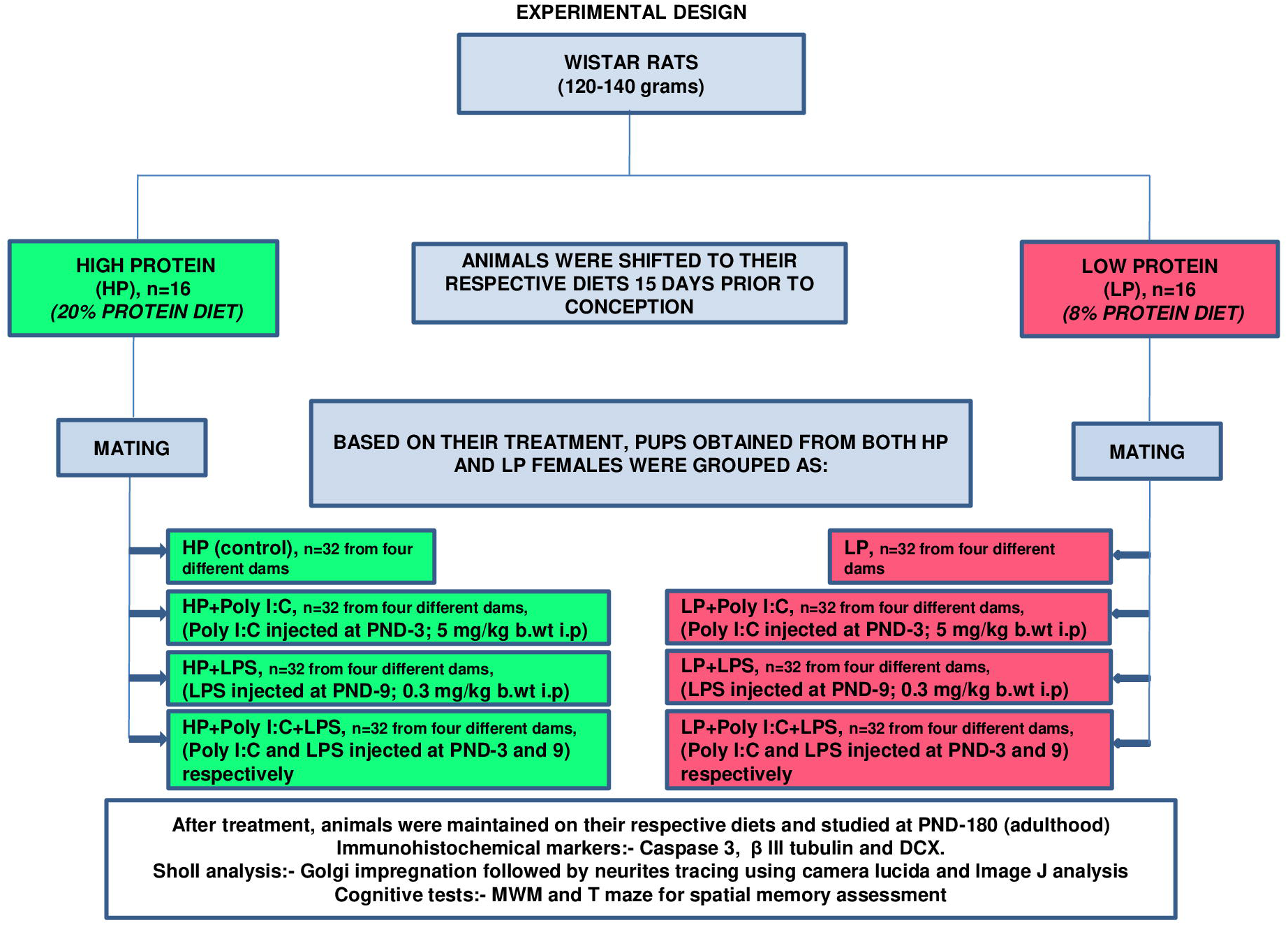
Representation of the experimental plan in a flow chart.

**Fig. 2.**
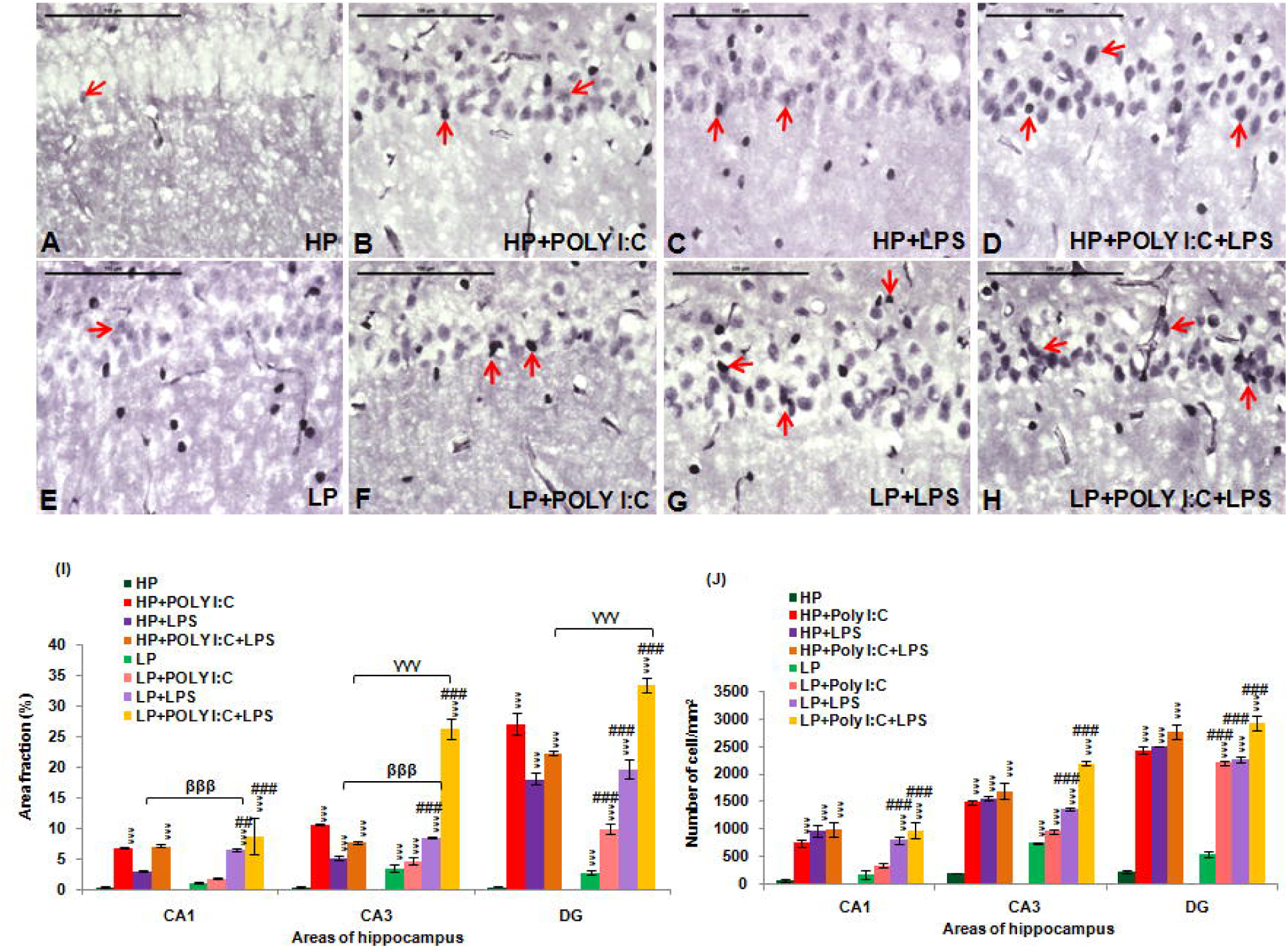
Active caspase 3 immunolabeled photographs and quantitative analysis of hippocampal sections demonstrating over-expression of activated caspase 3 protein at PND 180 following multi-hit stress: Multi-hit group i.e., LP+Poly I:C+LPS (H) presented maximum activated caspase 3 activity in comparison to control (A) and rest of the HP and LP treated groups (B, C, D, E, F, G) (red arrows). When LP and HP animals were subjected to single dose of either Poly I:C or LPS, the caspase 3 expression in LP+Poly I:C (F), LP+LPS (G), HP+Poly I:C (B) and HP+LPS (C) group was found to be higher than HP and LP alone group. (n=6 slides from different animals/group, scale bar=100μm) The quantitative analysis of active caspase 3 data (I) also shows maximum upregulation of caspase 3 protein in various hippocampal regions of LP+Poly I:C+LPS animals when compared to control and other treated groups. On administration of single stressor i.e., either Poly I:C or LPS to both HP and LP animals, HP+Poly I:C, HP+LPS, LP+PolyI:C and LP+LPS animalsshowed more activation of caspase 3 protein in the different regions of hippocampus when compared to HP and LP alone animals. Cell count graph (J), represents all the active caspase 3 positive cells in different groups. All the regions of hippocampus studied, showed same trend with LP+Poly I:C+LPS group having maximum number of cells, followed by HP+Poly I:C+LPS. Poly I:C and LPS treated HP and LP group also showed increased active caspase 3 positive cells, when compared to HP control and LP alone groups.(n=108 images each area from different slides/group). Values of One and Two Way ANOVA are expressed as mean±SEM; *P≤0.05, ***P≤0.001 with respect to controls; ^βββ^P≤0.001 with respect to HP+LPS and LP +LPS;^γγγ^P≤0.001 with respect to HP+Poly I:C+LPS and LP+Poly I:C+LPS and ^##^P≤0.005, ^###^P≤0.001 with respect to LP alone group

The changes vide supra were further confirmed through the quantitative data (Fig. 2I) with the area fraction of active caspase 3 immunopositivity being highly significant in the LP+Poly I:C+LPS group in the hippocampal regions (CA1, CA3 and DG) when compared with HP control (F_(7,856)_ =7.68, P≤0.001; F_(7,856)_ =13.97, P≤0.001; F_(7,856)_ =25.92, P≤0.001, group wise interaction) and LP alone group (F_(3,428)_ =6.92, P≤0.001; F_(3,428)_ =5.3, P=0.001; F_(3,428)_ =20.86, P≤0.001, interaction within treatments). A significant upregulation of caspase 3 in HP and LP animals following single-hit of either Poly I:C or LPS was also observed when compared with HP control and LP alone groups i.e., HP+Poly I:C in CA1 (F_(3,428)_ =5.98, P≤0.001), CA3 (F_(3,428)_ =29.25, P≤0.001) and DG (F_(3,428)_ =23.44, P≤0.001), LP+Poly I:C in CA3 (F_(7,856)_ =12.225, P≤0.001) and DG (F_(7,856)_ =8.445, P≤0.001; F_(3,428)_ =6.39, P≤0.001), HP+LPS in CA3 (F_(3,428)_ =13.608, P≤0.001) and DG (F_(3,428)_ =15.516, P≤0.001) and LP+LPS in CA1 (F_(7,856)_ =5.746, P≤0.001; F_(3,428)_=4.97, P=0.013), CA3 (F_(7,856)_ =22.92, P≤0.001; F_(3,428)_=14.19, P≤0.001) and DG (F_(7,856)_ =16.998, P≤0.001; F_(3,428)_=14.83, P≤0.001) respectively. Also, impact of LP diet was observed as on LPS and Poly I:C+LPS exposure, the LP animals reacted more vigorously than the corresponding HP groups and significant differences were found between HP vs. LP in CA3 (F_(1,642)_=2.93, P=0.038) and DG (F_(1,642)_=3.9, P≤0.001), HP+LPS vs. LP+LPS in CA1 (F_(1,642)_=3.26, P=0.021), CA3 (F_(1,642)_=9.36, P≤0.001) and HP+Poly I:C+LPS vs. LP+Poly I:C+LPS in CA3 (F_(1,642)_=6.8, P≤0.001) and DG (F_(1,642)_=3.6, P≤0.009).

While the area fraction data gave an idea of the intensity of labeling of active caspase 3 protein in the cells, total number of cells expressing active caspase 3 and going through apoptosis weredetermined by counting the number of active caspase 3 positive cells. From the cell count data (Fig. 2J), it was seen that Poly I:C, LPS and Poly I:C+LPS treatment to both HP and LP animals, hyped the number of cells expressing the active caspase 3 protein, therefore increasing the number of apoptotic cells in different hippocampal regions (CA1, CA3 and DG). Also, combinedexposure of Poly I:C and LPS to both HP and LP animals, led to maximum increase in number of apoptotic cells when compared to rest of the groups. Significant difference was found between HP and HP+Poly I:C (F_(3,428)_ =7.56, P≤0.001; F_(3,428)_ =18.2, P≤0.001; F_(3,428)_ =29.3, P≤0.001), HP and HP+LPS (F_(3,428)_ =9.9, P≤0.001; F_(3,428)_ =22.75, P≤0.001; F_(3,428)_ =30.1, P≤0.001), HP and HP+Poly I:C+LPS (F_(3,428)_ =10.36, P≤0.001; F_(3,428)_ =25.1, P≤0.001; F_(3,428)_ =33.8, P≤0.001), HP and LP+LPS (F_(7,856)_ =19.63, P≤0.001; F_(7,856)_ =19.63, P≤0.001; F_(7,856)_ =19.2, P≤0.001), HP and LP+Poly I:C+LPS (F_(7,856)_ =10.1, P≤0.001; F_(7,856)_ =33.6, P≤0.001; F_(7,856)_ =35.8, P≤0.001) in CA1, CA3 and DG whereassignificant difference between HP and LP (F_(1,642)_ =9.2, P≤0.001, impact of diet) was found in CA3 region and significant difference between HP and LP+Poly I:C (F_(7,856)_ =12.8, P≤0.001; F_(7,856)_ =14.2, P≤0.001), was found in CA3 and DG regions respectively. Within the LP groups also, there was significant difference in active caspase 3 positive cell number between LP and LP+Poly I:C in DG (F_(3,428)_ =10.04, P≤0.001), LP and LP+LPS (F_(3,428)_ =5.935, P≤0.001; F_(3,428)_ =10.34, P≤0.001; F_(3,428)_ =15.06, P≤0.001) and LP and LP+Poly I:C+LPS (F_(3,428)_ =9.01, P≤0.001; F_(3,428)_ =24.3, P≤0.001; F_(3,428)_ =15.06, P≤0.001) in CA1, CA3 and DG respectively.

### Catastrophe and downregulation of β III tubulin proteins resulted in fragmentation and loss of dendritic extensions

In the HP control animals, at PND180, the pyramidal neurons were morphologically preserved with apical dendrites, being evenly distributed and oriented in specific directions, devoid of any breakage or fragmentation (green arrows). Thecompactness of CA layer was also maintained in HP control animals (Figs. 3A).Whereas, in LP+Poly I:C+LPS i.e., multi-hit group the orientation of dendritic arbor was disrupted with void areas spottedin between crooked dendritic branches (Figs. 3H, red arrows). Additionally, the pyramidal cell layer was comparatively less dense with structurally altered and damaged neuronal population showing catastrophe of β III tubulin subunits when compared with HP control and similarly treated HP group (Fig. 3D). However, the LP alone group animals also presented a comparatively thin population of dendrites with mild β III tubulin labeling (Figs. 3E) as compared to the HP control group.Single-hit of Poly I:C or LPS also brought changes in the architecture of neurons in both HP (Figs. 3B, C) and LP animals (Figs.3F, G), which however was less than the multi-hit group.

**Fig. 3.**
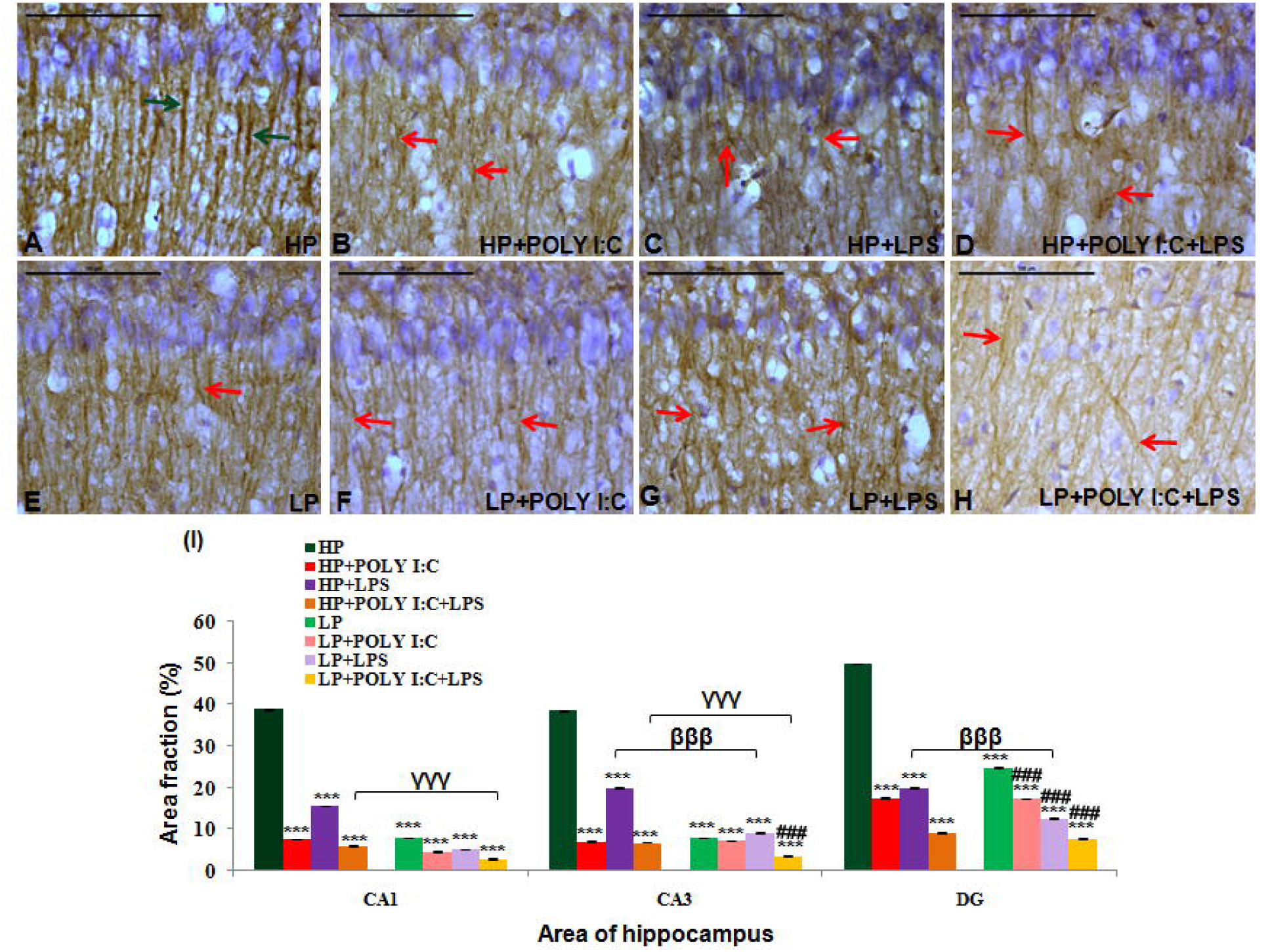
Light microscope images of β III tubulin labeled hippocampal CA regionalong with quantification data showing disorientation and misalignment of pyramidal neurons following multi-hit: Systematically orientated and intact neurons with long and continuous dendrites were spotted in adult HP controls (A, green arrows) whereas LP animals on combined exposure of both Poly I:C and LPS (LP+Poly I:C+LPS) showed incorrect neuronal orientation with void spacesin CA region. Their CA layers were loosely packed with tremendous β III tubulin catastrophe (H, red arrows). LP alone (E) and remaining stressed groups (HP+Poly I:C, HP+LPS, LP+Poly I:C, LP+LPS) showed some extent of dendritic damage with LP animals reacting more vigorously to stress when compared to their corresponding HP animals (B, C, D, F, G), (n=6 slides from different animals/group, scale bar=100 µm) The quantitative analysis of β III tubulin (I), showed significant decrease in density of β III tubulin in all the three regions of the hippocampus, following multi-hit exposure (LP+Poly I:C+LPS). The remaining LP and HP groups on Poly I:C and LPS treatment also had disintegrated microtubules with the LP animals losing β III tubulin more easily than the similar treated HP animals (n=108 images each area from different slides/group).Values of One and Two Way ANOVA are expressed as mean±SEM; ***P≤0.001 with respect to controls; ^βββ^P≤0.001 with respect to HP+LPS and LP +LPS, ^γγγ^P≤0.001 with respect to HP+Poly I:C+LPS and LP+Poly I:C+LPS and ^###^P≤0.001 with respect to LP alone group

The quantification data of β III tubulin mean area fraction (Fig. 3I) revealed a significant downregulation of β III tubulin following multiple stress with Poly I:C and LPS to LP animals in the sub regions of hippocampus studied i.e., CA1 (F_(7,856)_ =324.5, P≤0.001), CA3 (F_(7,856)_ =65.3, P≤0.001; F_(3,428)_=6.63; P≤0.001) and DG (F_(7,856)_ =263.2, P≤0.001; F_(3,428)_=6.09; P≤0.001) when compared with HP control (group wise interaction) and LP (interaction within treatments) alone group respectively (Fig. 3i).Single-hit with either Poly I:C or LPS to both HP and LP animals also led to a significant decrease in β III tubulin density in CA1, CA3 and DG regions when compared with HP control i.e., HP+Poly I:C (F_(3,428)_=204.63, P≤0.001; F_(3,428)_=34.2; P≤0.001; F_(3,428)_=211.3; P≤0.001), HP+LPS (F_(3,428)_=202.5, p≤0.001; F_(3,428)_=33.7, p≤0.001; F_(3,428)_=161.6, p≤0.001), LP+Poly I:C (F_(7,856)_=252.17; P≤0.001; F_(7,856)_=60.5; P≤0.001; F_(7,856)_ =252.84, P≤0.001) and LP+LPS (F_(7,856)_=221.16; P≤0.001; F_(7,856)_=63.9; P=0.002; F_(7,856)_ =252.8, P≤0.001). Also dystrophy of β III tubulin increased in LP+Poly I:C and LP+LPS groups mainly in DG region (F_(3,428)_ =5.34, P≤0.001, F_(3,428)_ =5.12, P≤0.001) when compared with LP alone group respectively. Impact of LP diet can be seen as significant differences were also observed within the HP and LP alone group in CA1 (F_(1,642)_=8.98, P≤0.001), CA3 (F_(1,642)_=11.81, P≤0.001) and DG (F_(1,642)_=7.165, P≤0.001); HP+LPS and LP+LPS group in CA3 (F_(1,642)_=6.06, P≤0.001) and DG (F_(1,642)_=4.2, P=0.003);HP+Poly I:C+LPS and LP+Poly I:C+LPS group in CA1 (F_(1,642)_=3.37, P=0.017) and CA3 (F_(1,642)_=6.14, P≤0.001) regions, showing vulnerability of LP animals on stress exposure.

### Early life stress led to decreased dendritic arborization, and dendritic length leading to a compromised neuronal profile, as analyzed through Golgi impregnation followed by Sholl analysis

Analysis of Golgi impregnated sections from various groups demonstrated the altered morphology of hippocampal pyramidal neurons on stress exposure. Fragmented dendrites as seen in β III tubulin labeled images were also observed in images of adult rat pyramidal neurons following single and multi-hit exposures. Acute stress generated in HP animals with single dose of either Poly I:C or LPS led to stunting of neurons, decrease in dendritic branches, drooping and haphazardly arranged dendritic arbors in HP+Poly I:C and HP+LPS animals (Figs. 4C b, c, Eb, c) when compared with HP control (Figs. 4A b c). Whereas chronic stress due to prolonged exposure of LP diet or combined administration of Poly I:C and LPS to LP or HP animals led to an abnormal extension of dendrites which however were fragile, incorrectly oriented and drooping (Figs. 4Bb, c, Db, c, Fb, c, Gb, C, Hb, c). Additionally, from the low magnified images of hippocampus, it was visible that on chronic stress exposure i.e., in LP+Poly I:C+LPS animals, the CA layers became improvident due to increased density and overcrowding of damaged dendrites (Figs. 4H a).

**Fig. 4.**
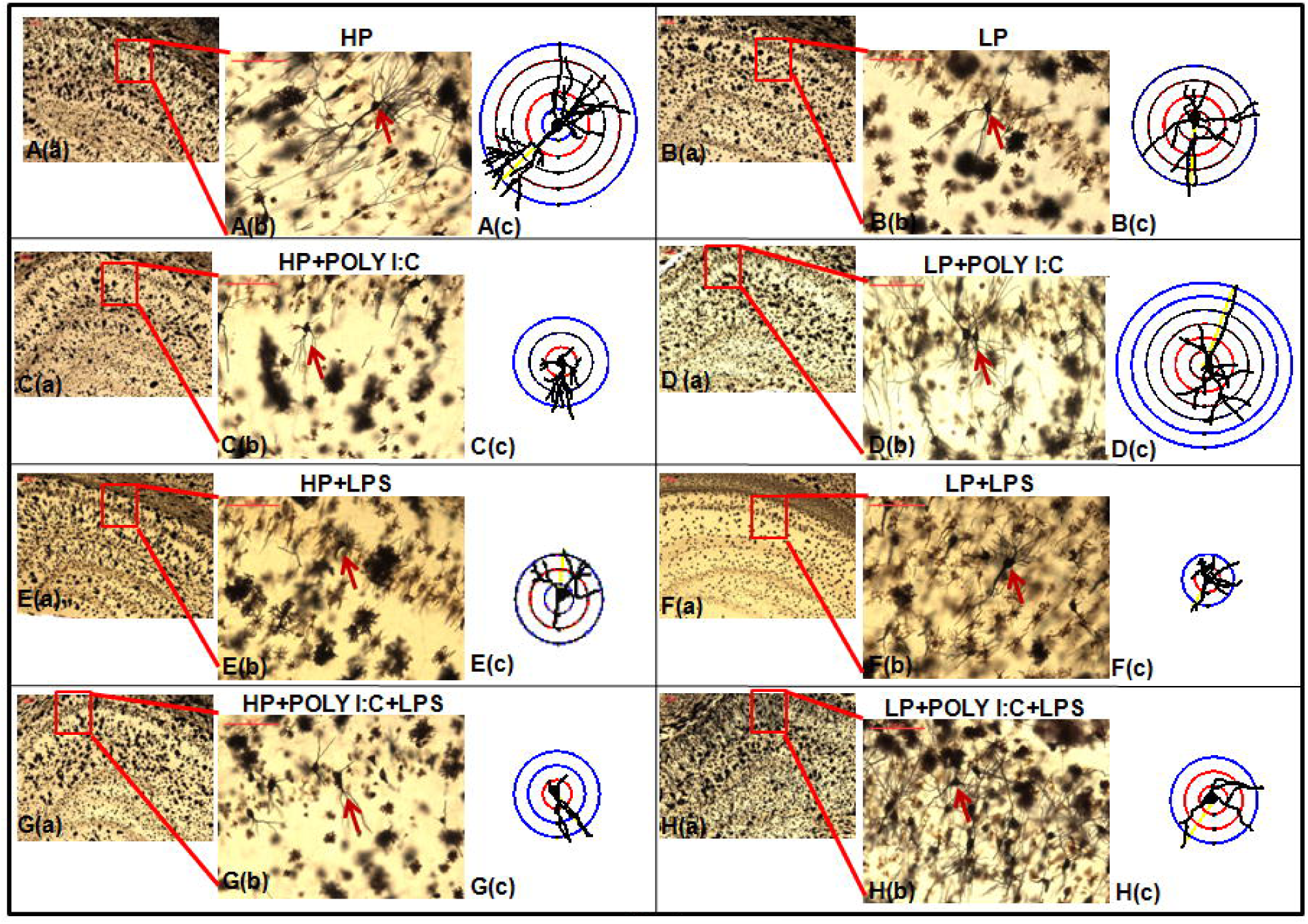
Microscopic as well as camera lucida traced Golgi impregnated images of neurons showing abnormal neuronal morphology on stress exposure: Morphological damage including loss of dendritic branching leading to neuronal stunting and dendritic drooping was seen in chronically stressed LP+Poly I:C+LPS group (Ha, b, c), when compared with morphologically intact neurons belonging to HP controls (Aa, b, c). On single-hit with either Poly I:C or LPS, two type of morphological alteration was observed in both HP and LP animals. Some neurons on acute stress were found to be stunted (Ca, b, c, Ea, b, c, Fa, b, c, Ga, b, c) while others due to chronicity of stress, were observed to compensate for the damage by extending their primary dendrites (Da, b, c, Ba, b, c), (n=6 slides from different animals/group, scale bar=100μm)

Sholl analysis on camera lucida traced neuronal images was performed to assess morphometric variation including dendritic arborization and length among treated and control groups. Total number of Sholl circles at an interval of 20µm gave length of dendrites whereas dendritic arborization was calculated from the mean total number of intersections made by dendrites on the circles.Specifically, pyramidal neurons were chosen for analysis and hence as per the shape of such neurons, numbers of intersecting points in the Sholl circles were maximum at 40µm, which further decreased with increase in the length of the neurons. The histogram was plotted with values of the distance from the soma by X axis and mean number of intersections between dendrites and Sholl circles by Y axis, which gave an idea of the morphometric differences between pyramidal neurons of HP control and other treated groups (Fig. 5). The arborization of dendrites was decreased in LP alone animals, with average dendritic length being confined till 100μm, when compared with 180μm long dendritespersistently seen in HP animals. On individual Poly I:C and LPS treatment, the dendritic arborization and length further decreased in HP and LP animals i.e., HP+Poly I:C (140μm), HP+LPS (60μm) and LP+LPS (60µm). Compensatory increase in dendritic length was observed in LP+Poly I:C animals when compared with LP alone group as the average length of dendrites of LP+Poly I:C group was observed to be 140µm long. On combined exposure of Poly I:C and LPS to HP and LP animals i.e., in HP+Poly I:C+LPS and LP+Poly I:C+LPS groups, the average dendritic length and arborization further declined when compared to HP control and LP alone animals. (Fig. 5). (The statistical data for comparison of mean arborization is shown in Table 1, given under supplementary data).

**Fig. 5.**
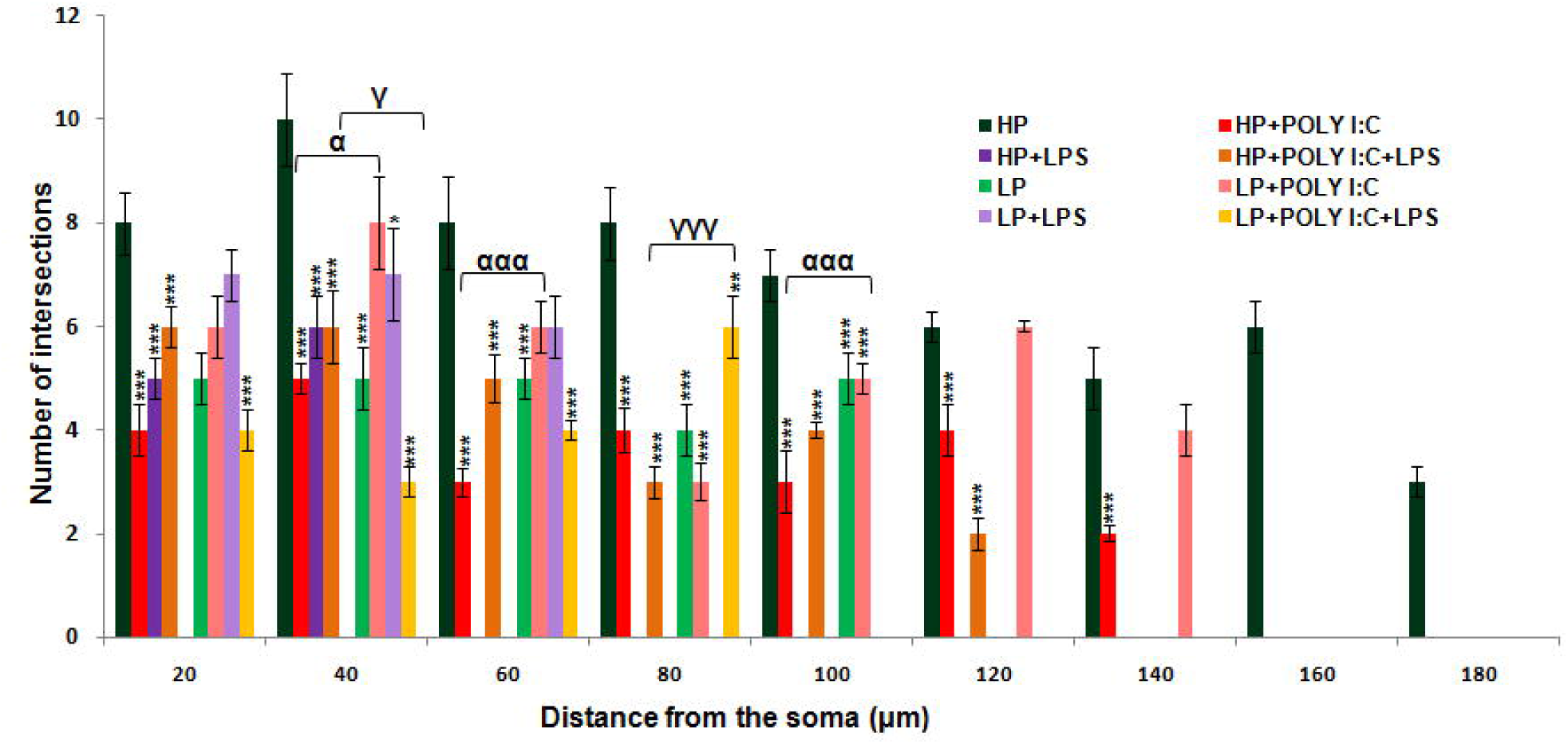
The histogram shows significant difference in length and arborization between HP control and treated groups on Sholl analysis: The length of the pyramidal neurons denoted as distance from the soma by X axis were affected in all the treated groups. Arborization, denoted as the number of intersections by Y axis were also found to be decreased among treated groups, with minimum arborization seen in LP+Poly I:C+LPS group. (n=108 neurons from 6 different slides/group).Values of One and Two Way ANOVA are expressed as mean±SEM; *P≤0.05, **P≤0.005, ***P≤0.001 with respect to controls; ^α^P≤0.05, ^ααα^P≤0.001 with respect to HP+Poly I:C and LP+Poly I:C; ^γ^P≤0.05, ^γγγ^P≤0.001 with respect to HP+Poly I:C+LPS and LP+Poly I:C+LPS

### Stress triggered spotting of immature neurons, identified by anti-DCX immunolabelling of naive neurons

Damage induced increase in DCX positive neurons (arrows), was observed in treated groups. The spotting of DCX positive neurons in CA layer were directly proportional to the amount of damage that occurred to residential neurons. Chronic stress due to multiple-hitin LP animals i.e., LP+Poly I:C+LPS (Figs. 6H, 7H) led to a vigorous increase in DCXpositive cells in and around CA layers and DG. Additionally, the DCX positive cells were not discrete and appeared to be clumped in DG may be because of chronic stress related damage to naive DG neurons. Comparatively, no such hype in DCX positive cells were seen in HP+Poly I:C+LPS group (Figs. 6D, 7D), which could be due to the ability of HP animals to rectify and compensate for chronic immune activated neuronal damage by incorporation of the naive neurons into the circuitry. HP control animals had very few numbers of DCX positive neurons in the CA layers (Fig. 6A) but prominent DCX cells were present in the DG notch suggesting healthy adult neurogenesis (Fig. 7A). On the other hand, LP alone group was seen to have more DCX expressing cells in the CA layers but however in the DG, less DCX positive cells were visible (Figs. 6E, 7E). The single exposure of Poly I:C tobothHP and LP animals also led to a hype in DCX positive cells in both CA layer and DG (Figs. 6B,F, 7B,F). No such prominent hype was seen in LPS exposed HP and LP groups (Figs. 6C,G, 7C,G).

**Fig. 6.**
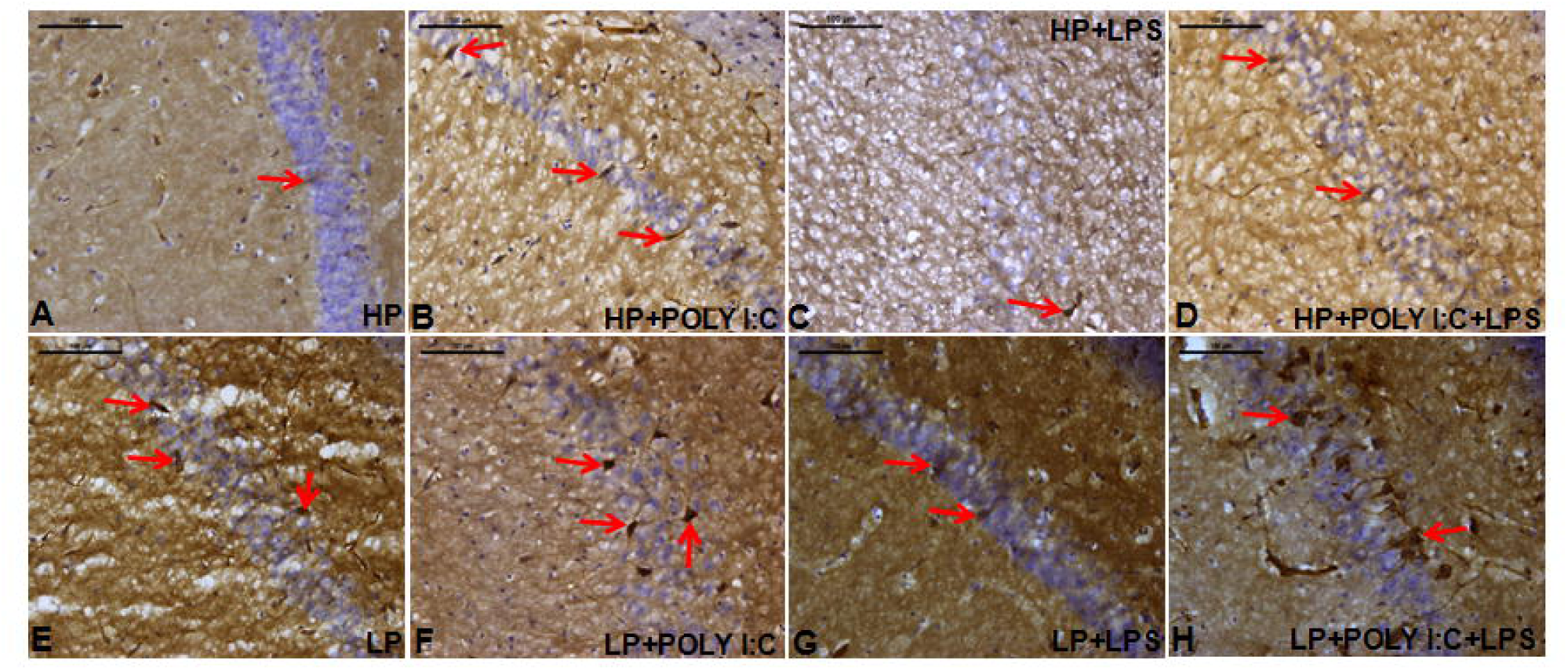
Microscopic images of hippocampus showing stress dependent increase in DCX expression: Stress induced increase in DCX expressing cells were observed in LP+Poly I:C+LPS group with cells clumped around CA layer (H, red arrows). Poly I:C and LPS were also found to increase DCX expression in both HP and LP animals (B, C, D, F, G), (red arrows), when compared with HP (A) and LP (E) alone group but again was less than the LP+Poly I:C+LPS group. (n=6 slides from different animals/group, scale bar=100µm)

**Fig. 7.**
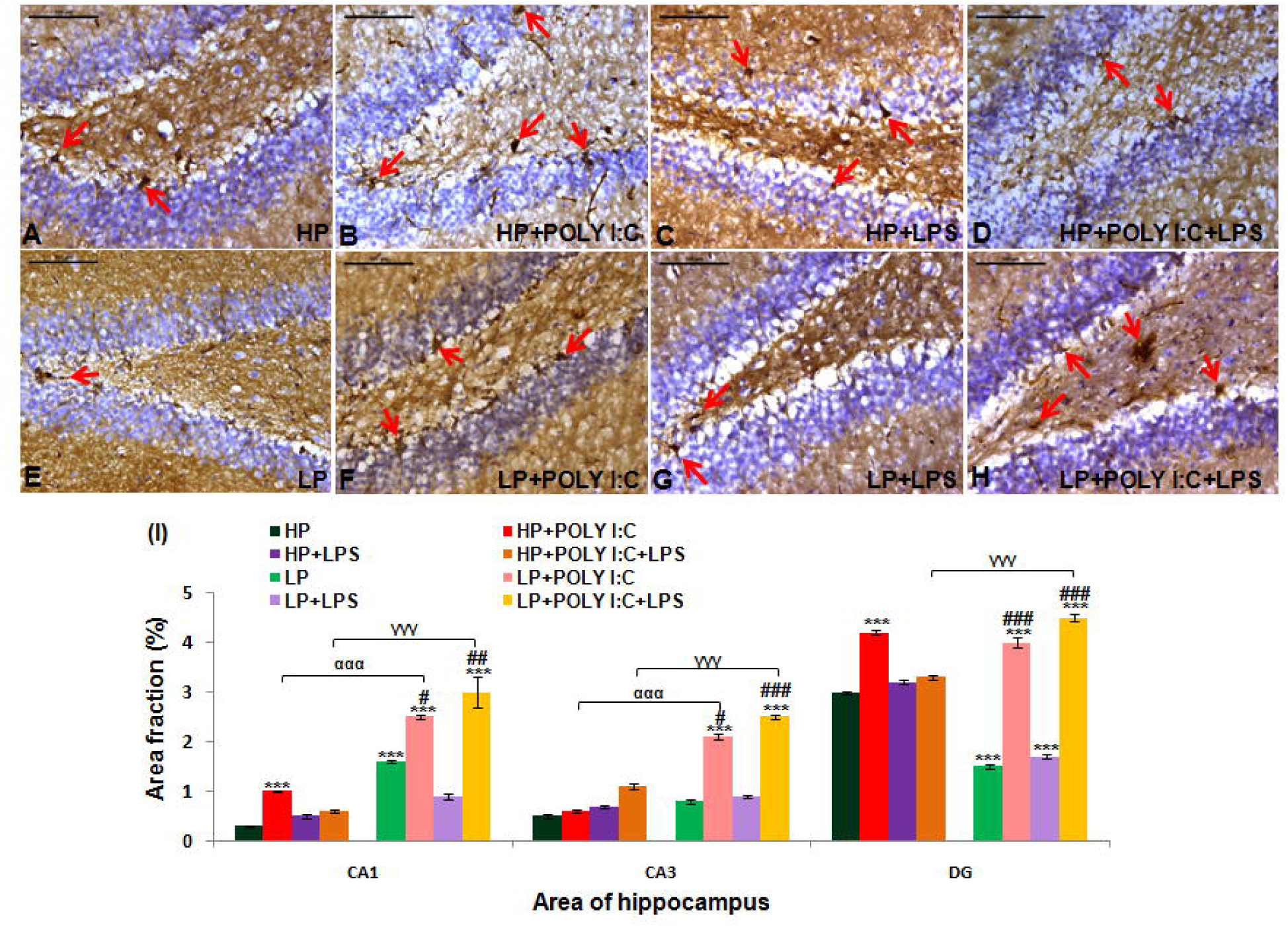
The photographs of DG and quantification data of DCX protein, showing upregulation of DCX in the multi-hit group: The spotting of DCX positive cells were heightened on combined exposure of Poly I:C and LPS in both HP and LP animals (D, H, red arrows), showing moreDCX expressing new neurons when compared with HP (A) and remaining treated groups (B, C, E, F). (n=6 slides from different animals/group, scale bar=100µm) The graphical representation of mean area fraction (%) of DCX proteins (I) also demonstrated that LP+Poly I:C+LPS group had highest expression of DCX proteins in all the regions of the hippocampus when compared to the other treated groups. (n=108images each area from different slides/group).Values of One and Two Way ANOVA are expressed as mean±SEM; ***P≤0.001 with respect to controls; ^ααα^P≤0.001 with respect to HP+Poly I:C and LP+Poly I:C; ^γγγ^P≤0.001 with respect to HP+Poly I:C+LPS and LP+Poly I:C+LPS and ^#^P≤0.05, ^##^P≤0.005, ^###^P≤0.001 with respect to LP alone group

The morphometric analysis of DCX immunoreactivity in CA1, CA3 and DG (Fig. 7I) revealed highest upregulation of DCX protein in LP+Poly I:C+LPS group when compared to HP control (F_(7,856)_ =7.83, P≤0.001; F_(7,856)_ =45.93, P≤0.001; F_(7,856)_ =88.74, P≤0.001), LP alone (F_(3,428)_ =4.89, P=0.003; F_(3,428)_ =6.76, P≤0.001) and HP+Poly I:C+LPS groups (F_(1,642)_ =8.06, P≤0.001; F_(1,642)_ =5.78, P≤0.001; F_(1,642)_ =8.34, P≤0.001). Such increase in DCX immunopositivity was also seen inLP+Poly I:Cgroup when compared with HP control and LP alone group in CA1, CA3 and DG (F_(7,856)_ =28.3, P≤0.001; F_(7,856)_ =15.9, P≤0.001; F_(7,856)_ =28.7, P≤0.001, F_(3,428)_ =4.183, P=0.016; F_(3,428)_ =4.79, P=0.004; F_(3,428)_ =4.009, P=0.02) and in HP+Poly I:C groupwhen compared to HP control in CA1 and DG(F_(3,428)_ =16.8, P≤0.001; F_(3,428)_ =57.7, P≤0.001) respectively.Within the group significant difference were found between HP control vs. LP alone group in CA1 and DG (F_(1,642)_ =16.5, P≤0.001; F_(1,642)_ =47.3, P≤0.001), HP+Poly I:C vs. LP+Poly I:C in CA1 and CA3 (F_(1,642)_ =26.5, P≤0.001; F_(1,642)_ =37.3, P≤0.001) and HP+Poly I:C+LPS vs. LP+Poly I:C+LPS in CA1, CA3 and DG(F_(1,642)_ =27.83, P≤0.001; F_(1,642)_ =25.93, P≤0.001; F_(1,642)_ =18.74, P≤0.001).

### Impaired spatial memory during MWM and T maze task, on exposure to various early life stressors MWM

Animals were trained to use spatial clues for navigation and localization of escape platform during MWM task. From the latency and mean path efficiency graph (Fig. 8I, J), it was noted that due to severe chronicity, the latency was highest and path efficiency was lowest in multi-hit, i.e., LP+Poly I:C+LPS animals when compared with HP control (F_(7,88)_ =16.17, P≤0.001, F_(7,88)_ =11.4, P≤0.001) and LP alone group (F_(7,88)_ =13.59, P≤0.001), which meant that they took more time and followed complex path while reaching the escape platform. LP alone animals were also found to have an increasedlatency and decreased path efficiency depicting impaired spatial memory when compared to HP control (F_(1,66)_=9.54; P≤0.001, impact of LP diet) which on further immune activation with either Poly I:C or LPS was heightened as the latency increased and path efficiency decreased in LP+Poly I:C and LP+LPS animals when compared to HP control (F_(7,88)_=5.6; P≤0.001; F_(7,88)_=5.8; P≤0.001) and similarly treated HP groups i.e., HP+Poly I:C (F_(1,66)_=5.9; P≤0.001) group. Significant difference in latency were also found between LP, LP+Poly I:C (F_(3,44)_ =10.4, P≤0.001) and LP, LP+LPS groups (F_(3,44)_ =12.6, P≤0.001). Also, when percent time spent in target quadrant data was analyzed (Fig. 8K), it was seen that HP control group had maximum time spent in the target quadrant to which the platform belongs, avoiding the other zones. HP+Poly I:C, HP+LPS and HP+Poly I:C+LPS had reduced percent time spent denoting poor memory and incapability of the treated groups to spot the quadrant containing platform. Significant difference was found between HP and HP+Poly I:C+LPS group (F_(3,44)_=8.25; P≤0.001). LP alone group had low percent time spent when compared to HP control (F_(1,66)_=7.01; P≤0.001), which further reduced on Poly I:C, LPS and Poly I:C+LPS exposure (F_(3,44)_=7.2; P≤0.001, F_(3,44)_=7.9; P≤0.001, F_(3,44)_=9.4; P≤0.001) when compared to HP control group.Thus, from the above-mentioned data it can be seen that combo exposure acted synergistically with LP treatment and extravagated the damage in LP animals. However, such effects of multi-hit stress were much less severe in HP animals may be because of their ability to resist or rectify.

**Fig. 8.**
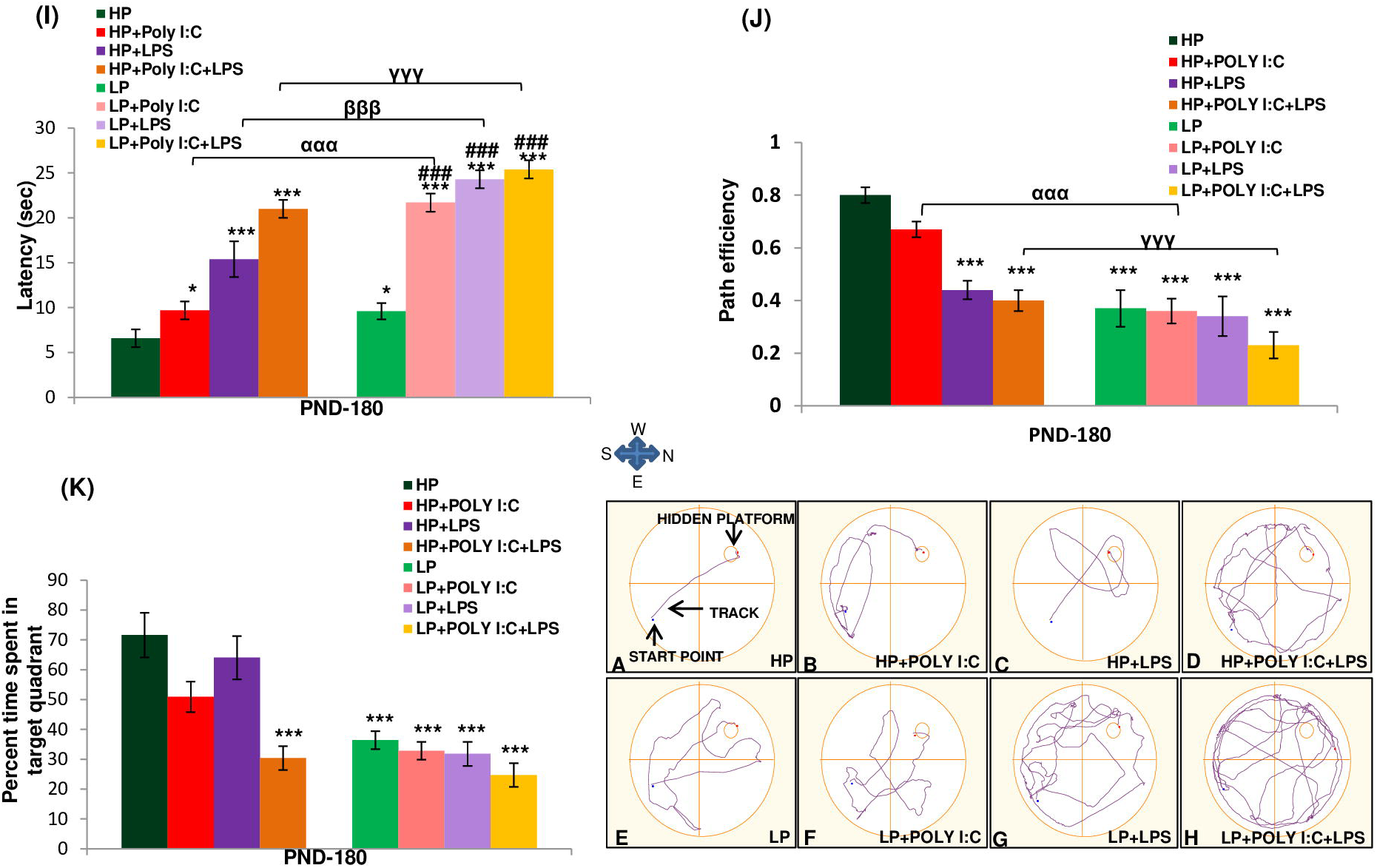
MWM data showing memory impairment in rats on stress exposure: The track records of animals during MWM task demonstrates maximum memory impairment in LP+Poly I:C+LPS group (H) when compared to HP control (A) and other treated groups (B, C, D, E, F, G). The LP animals on treatment with either Poly I:C or LPS (F, G) also performed poorly than the HP control (A) and corresponding HP treated group (B, C), (n=12/group). The graphical analysis of mean latency, path efficiency and percent time spent in target quadrant (I, J, K) depicts memory impairment in HP and LP animals on Poly I:C and LPS administration as the animals belonging to treated groupstook longer time to reach the escape platform, followed complicated trajectory and spent less time in the target quadrant. Maximum deficit was shown by LP+Poly I:C+LPS animals, when compared to rest of the groups. Within the group significant difference was found between HP+Poly I:C and LP+Poly I:C animals, showing that on administration of Poly I:C or LPS, LP animals suffered more damage when compared to HP animals. (n=12/group).Values of One and Two Way ANOVA are expressed as mean±SEM; ***P≤0.001 with respect to control; ^ααα^P≤0.001 with respect to HP+Poly I:C and LP+Poly I:C; ^γγγ^P≤0.001 with respect to HP+Poly I:C+LPS and LP+Poly I:C+LPS

From the track records of MWM task, indirect and complex trajectory depicting impaired spatial memory was seen in LP (Fig. 8E) alone, LP+Poly I:C (Fig. 8F), and LP+LPS (Fig. 8G) groups. The HP control animals (Fig. 8A) followed a direct path from the start point to the escape platform which the treated animals failed to do. The complexity of tracks and impairment of memory increased in with increase in chronicity of stress and hence, the LP+Poly I:C+LPS animals as they took indirect and haphazard path and reached the escape platform after doing multiple errors when compared with other groups (Fig. 8H). Single or combo exposure of Poly I:C or LPS to HP animals also showed some impairment in path efficiency (Figs. 8B, C, D), but was comparatively lesser than the respective LP group animals.

### T Maze

Alternate baited arm protocol was followed in T maze task to check spatial working memory among all the groups. The mean path efficiency data was plotted in graph (Fig. 9I), from which, it was interpreted that spatial working memory was impaired following stress. Such spatial memory impairment was seen to be highest in multi-hit group i.e., LP+Poly I:C+LPS as the animals were not able to distinguish between left and right arms, they committed more error while choosing the correct arm to obtain reward (F_(7,88)_=9.3; P≤0.001). Similar to MWM, the LP alone animals performed less efficiently than HP controls (F_(1,66)_=5.53; P≤0.001). On additional treatment with Poly I:C or LPS, the path efficiency depicting spatial working memory further declined in LP animals when compared with HP control (F_(7,88)_=7.57; P≤0.001; F_(7,88)_=6.3; P≤0.001) and the impairment in LP+Poly I:C (F_(1,66)_=5.072; P≤0.001) and LP+LPS and LP+Poly I:C+LPS groups was more than their HP counterparts.

**Fig. 9.**
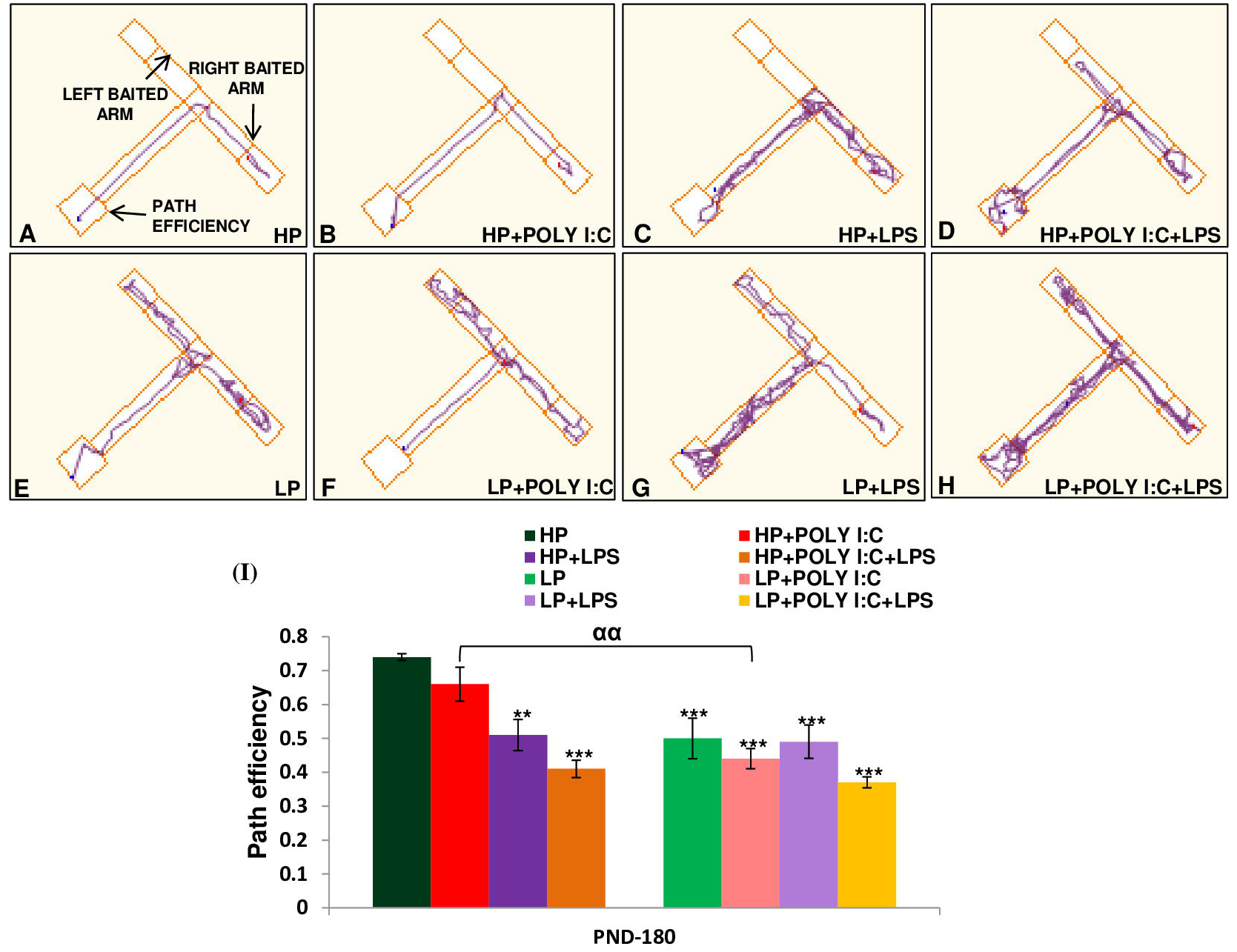
The mean path efficiency histograms and tracks during T maze task shows maximum decline in path efficiency indicating poor memory in LP+Poly I:C+LPS group: Unavailability of right left discrimination with poor path efficiency in LP+Poly I:C+LPS group (H) is visible from the track records of T maze task. LP alone treated animals also had complex and impaired tracks (E) which further deteriorated on Poly I:C or LPS treatment (F, G). HP animals on similar treatment with Poly I:C and LPS (B, C, D) also had impaired tracks when compared with the tracks of HP control (A) but however, the damage in tracks were comparatively lesser than the LP animals, (n=12/group). HP and LP treated Single-hit groups showed spatial memory impairment with decrease in path efficiency as seen from the T maze (I) when compared to HP control but damage was chronic in case of multi-hit group when Poly I:C and LPS was administered sequentially in LP animals i.e., LP+Poly I:C+LPS group. (n=12/group).Valuesof One- and Two-Way ANOVA are expressed as mean±SEM; **P≤0.005, ***P≤0.001 with respect to controls; ^αα^P≤0.005, with respect to HP+Poly I:C and LP+Poly I:C

The track records revealed distorted and repeated track in treated animals showing errors while choosing the right baited arm. It was evident from the complex and haphazard track of LP+Poly I:C+LPS animals that they had maximum impairment in their spatial working memory when compared to other treated groups (Fig. 9H). Moreover, LP (Fig. 9E), LP+Poly I:C (Fig. 9F) and LP+LPS (Fig. 9G) animals also had repeated tracks showing memory impairment when compared to HP control (Fig. 9A) and similarly treated HP groups (Figs. 9B, C, D). Lastly, with in the HP groups also, there was impairment in memory upon treatment with Poly I:C and LPS either singularly or in combination.

## Discussion

The role of perinatal stress as causative factor for neurological disorders like Schizophrenia, Alzheimer’s, Parkinson’s etc. has been well reported (Huang, 2014, Hoeijmakers et al., 2015, Hohmann et al., 2017). However, the mechanism of action of stressors and the influence of one type of stressor on another still remains unclear. Some of the studies have focused the dual-hit hypothesis according to which, first hit at genetic level during critical developmental periods increases the risk of second-hit, which is generally an environmental insult like infection and nutritional deficiency. Dual-hit exposure may act synergistically and give rise to Schizophrenic symptoms in adult individuals (Maynard et al., 2001; Feigenson et al., 2015). However, due to a wide range of stressors and their diverse mode of action, dual-hit hypothesis alone is not sufficient to provide a link between stressors and neurological disorders. This study is the first one to address and streamline the events that lead to symptoms seen during CNS malfunctioning in a multi-hit model.

In this study, perinatally multi-hit animals were found to have highest active caspase 3 expressions in hippocampus when compared with other groups and additionally, dendrites of such chronically stressed animals also showed active caspase 3 expressions depicting their degeneration and neuronal dysfunction. Such upregulation of active caspase 3 could be a reason for low β III expression in multi-hit groups. The multi-hit animals also subsequently exhibited damaged neuronal profile and decreased dendritic arborization, may be because of vigorous active caspase 3 triggered β III tubulin damage. Simultaneous upregulation of DCX protein was also found in DG and CA layers of multi-hit animals with chronic stress leading to scattering of DCX positive cells in CA layers. Such altered cytoarchitecture of neurons in hippocampus was finally reflected as the impaired spatial memory in stressed animals which was directly proportional to the chronicity of stress. Lastly, it was also seen that LP animals were prone to infection as they reacted more vigorously to a subsequent stress exposure when compared to HP animals suggesting thereby that the stressors are interdependent and act synergistically which has been variously hypothesized (Schaible and Kaufmann, 2007; Katona and Apte,2008).

Active caspase 3 is an apoptotic marker used to study the extent of cellular damage during stress exposure (Mcllwain et al., 2015). Chronic stress in multi-hit (LP+Poly I:C+LPS) animals was found to be responsible for increase in caspase 3 protein in hippocampus, which was higher than the single-hit groups. Such chronic stress dependent increase in active caspase 3 protein was also reported in a depressed rat model by Bachis and co-workers (2008). Restrained stress when combined with forced swim stress was also found to increase the caspase 3 level in prefrontal cortex of Wistar rats (Zhang et al., 2017). Moreover, Toll like receptors (TLR) activated by viral and bacterial pathogens along with oxidative stress are also found to be responsible for activating caspases and inducing apoptosis in cells leading to neurodegeneration (Snigdha et al., 2012; Loeches et al., 2014; Sharma et al., 2016). Active caspase 3 dependent neuronal death in case of specific virus-oriented encephalitis, viz., West Nile virus encephalitis and Japanese encephalitis are also being reported (Samuel et al., 2007; Mishra and Basu, 2008). Thus, all the single stress-oriented studies (vide supra) indicate that caspase 3 mediated cell death is a major contributor to neurodegeneration and hence tally with the active caspase 3 data presented in this study. Moreover, the combined exposure of multiple stressors (multi-hit) is responsible for vigorous cell death in hippocampus, further accelerating the occurrence of neurodegeneration and related disorders.

Beside cell death, activated caspase 3 also causes microtubule distortion leading to dendritic loss and stunting in neurons, prominent in Alzheimer’s disease (Troy and Jean, 2015). Stress dependent damage of mitochondrial membrane also leads to activation of caspase 3 in dendrites, causing fragmentation (Mattson et al., 1998; Amelio et al., 2011). Such activecaspase 3 labeled dendrites were also seen in chronically stressed multi-hit group in the present study confirming their fragmentation. Moreover, caspase 3 has been reported to cleave the tubulin protein causing both axonal and dendritic degeneration (Sokolowski et al., 2014). Thus, the caspase 3 activation during various stress conditions may be a crucial factor leading to β III tubulin degradation.

Multi-hit stress in our study led to an extensive β III tubulin degeneration in neurons, which could be linked to compromised dendritic arborization of neurons and decreased neuronal profiling observed in the multi-hit animals. Such alterations in microtubule dynamics is also observed in case of Alzheimer’s disease as microtubule destabilization is associated with formation of neurofibrillary tangles and aggregation of hyperphosphorylated tau (Brandt and Dakota, 2017). Microtubule instability is also common in other neurodegenerative disorders like Parkinson’s disease and Amyotrophic Lateral Sclerosis (Dubey et al., 2015; Salama et al., 2018). Thus, from our β III tubulin data it can be interpreted that multi-hit stress can make an individual prone to neurological disorders as cytoskeletal changes may lead to neurodegeneration leading to behavioral and cognitive deficits through compromised neuronal arbors.

Stunted neurons due to fragmented and drooping dendrites were common in chronically stressed multi-hit group. Although no multi-hit studies have been reported so far, our result can still be supported by the single-hit studies which reported that following viral or bacterial infections, the CA1 neurons in rats becomes architecturally damaged with reduced dendritic length and arborization further leading to cognitive impairment (Vyas et al., 2002; Radley et al., 2004; Murmu et al., 2006; Baharnoori et al., 2009; Jurgens et al., 2012). Thus, multi-hit stress could be a major factor that accelerates cellular damage further causing memory impairment leading to neurodegenerative disorders.

Alongside cellular damage, stressors are also reported to enhance adult neurogenesis in order to compensate for regional damage. In the present study as well, the DCX positive cells were spotted around CA layers and DG of stressed animals. DCX is reported to be strongly expressed by migrating immature neurons (Kaindl et al., 2006; Klempin et al., 2011; Madhyastha et al., 2013) and in our study, mainly after the multi-hit exposure, DCX expression was upregulated, suggesting an increase in migration of newly generated neurons. Some studies have reported the difference in extent of neurogenesis between sub ventricular (SVZ) and sub granular (SGZ) zone, stating that SVZ regions contains more neural stem cells and with additional amplifying signals, the extent of neurogenesis in SVZ is somewhat higher than SGZ region (Zhu et al., 2014; Ghosh et al., 2019). However, some studies have also reported a decrease in DCX positive cells in hippocampus of rats on chronic stress exposure (Dagyte et al., 2009). This could be because that DCX positive immature neurons were unable to be incorporated in to the circuitry due to chronicity of stress and hence were prone to degeneration suggesting that increase in chronicity decreases the chance of maturation and survival of neurons born as a result of compensation mechanism (Nacher et al., 2004, Lemaire et al., 2005; Naninck et al., 2014). Hence in our study even with an increase in adult neurogenesis, the cognitive abilities of stressed animals remain impaired with the multi-hit animals showing maximum memory impairment in MWM and T maze task which can be further linked with occurrence of neurodegeneration related disorders in adult animals.

Memory impairment is a hallmark of neurodegenerative disorders which occurs due to stress dependent altered neuronal circuitry (Esch et al., 2002; Akers et al., 2006; Wen et al., 2017). Early life exposure to stressors like Poly I:C, LPS and perinatal protein malnourishment individually are well reported to cause memory impairment in rats (Naik et al., 2015; Zakaria et al., 2017; Baghel et al., 2018). However, in the present study all these early life stressors in combination were found to heighten later life memory impairments. Such multi stress exposure during early life was found to cause chronic damage to spatial memory in LP+Poly I:C+LPS animals via stress dependent caspase 3 activation and β III tubulin catastrophe leading to loss of neuronal connectivity and subsequent neurodegeneration in adult rats.

## Materials and methods

### Animal husbandry and early life stress induction

Wistar rats were maintained in the animal house facility under controlled physical environment (temperature=25±1° C, humidity=65±2%, light and dark cycle=12 hr). Before shifting to the experimental diet, all F_0_ animals were given ad libitum access to clean drinkable water (reverse osmosis water) and rat pellet feed. 32 Virgin females (3 months old, body weight; b.wt., 120-140gms) were selected and shifted to control i.e., high protein (HP; 20 % protein, n=16) and low protein (LP; 8% protein, n=16) diets, 15 days prior to mating and then maintained on their respective diets throughout gestation and lactation periods (isocaloric rat feed, both 8% and 20% protein diet were obtained from National Institute of Nutrition, Hyderabad, India). The day of parturition was noted as postnatal day 0 (PND 0). The F_1_ pups from both HP and LP dams were used for creating the following groups. Litter size adjusted to 8 to avoid variation.

### Control

HP F_1_ pups (n=32, from four different dams) without any treatment were considered as controls. They were maintained with their respective dams and used for various studies at the age of PND 180 (b.wt. 230±10gms).

### Low protein model (LP, single stress)

F_1_ pups from LP dams (n=32, from four different dams) were used as LP alone group. The animals were maintained in LP diet and used for various studies at the age of PND 180 (b.wt. 180±10gms).

### Viral infected model in HP and LP groups (HP+Poly I:C and LP+Poly I:C)

Poly I:C (Sigma; St. Louis USA) was injected to mimic a viral infected model. Stock solution was prepared by dissolving 5mg of Poly I:C in pre-heated (60°C) TBE buffer (Tris-Borate-EDTA). The solution was mixed properly and stored at 4°C for further use. Equal number of pups (n=32, from four different HP as well as LP dams) were injected with Poly I:C (IP) at PND 3 at a dose of 5mg/kg b.wt. The pups were then returned to their respective dams and maintained till used at PND 180 (HP+Poly I:C, b.wt. 230±10gms and LP+Poly I:C, b.wt. 180±10gms) according to the experimental plan.

### Bacterial infected model in both HP and LP groups (HP+LPS and LP+LPS)

LPS (Sigma Aldrich, *E. coli*, serotype 0111:B4), was prepared by dissolving 0.3 mg of LPS in 1ml of PBS (Phosphate buffer saline). Similar to Poly I:C treatment, equal number of pups from both HP and LP dams (n=32 from four different dams) were injected with LPS (IP) at PND 9 at a dose of 0.3 mg/kg b. wt and then returned to their respective dams and maintained till used accordingly at PND 180 (HP+LPS; b.wt. 230±10gms and LP+LPS; b.wt. 180±10gms).

### Viral and bacterial combined infected (multi-hit) model in both HP and LP groups (HP+Poly I:C+LPS and LP+Poly I:C+LPS)

Pups from both HP and LP groups (n=32 from four different dams) were injected (IP) with both Poly I:C and LPS at the same dose and postnatal days similar to the viral and bacterial infected HP and LP groups (PND 3 and 9, Dose-5mg/kg and 0.3 mg/kg b. wt., respectively). The animals were then used for various studies at PND 180 (HP+Poly I:C+LPS, b.wt.:230±10gms and LP+PolyI:C+LPS; b.wt., 180±10gms).

Equal number of male and female rats was used in each experiment as post hoc analysis did not bring out any sex specific differences and the data from both the sexes was combined together and presented as average values. Controls were injected with vehicle alone in correspondence to Poly I:C and LPS exposure. For accuracy, all injecting procedures were performed using Stoelting Nanoinjector and Hamilton microsyringe under aseptic conditions. Litter size of 8-9 pups per dam was used to avoid any variation. The overall study consists of 8 groups, 4 from control diet (HP, HP+Poly I:C, HP+LPS, HP+Poly I:C+LPS) and 4 from LP diet (LP, LP+Poly I:C, LP+LPS, LP+Poly I:C+LPS) and are detailed in Fig. 1.

### Perfusion and tissue harvesting for immunocytochemistry

Animals from all the above-mentioned groups (n=6/group, from different dams) wereanesthetized using diethyl ether and transcardially perfused at PND 180, using pre chilled PBS (phosphate buffer saline, 0.01 M, pH-7.4) followed by 2% paraformaldehyde prepared in 0.01 M PB (Phosphate buffer). The brains were carefully dissected out and then post fixed for 24 hrs in same fixative, using immersion fixation technique. The tissues were cryoprotected in sucrose gradients (10%, 20%, 30% sucrose in PB) for consecutive days. The tissues were then sectioned through occipito-temporal region containing hippocampus (14 µm thickness) using a cryotome machine (Leica CM1900, Germany) followed by storage at -20°C for immunohistochemical analysis.

### Immunolabelling of active caspase 3 protein using immunoenzymatic method with nickel enhancement

Brain sections from each group (n=6 slides, from different animals and dams/group) were carefully selected and air dried followed by TBS (0.05 M Tris Buffered Saline, pH 7.4) washing. Membrane permeabilization was done for 20 minutes using 0.5% Triton X-100 (Sigma) in TBS. The sections were then washed with TBS, blocked in 1% H_2_O_2_ (Merck) and incubated with 1% NGS (Normal Goat Serum, Vector PK6200) for 90 minutes. After non-specific protein blocking with NGS, the sections were incubated overnight at 4°C with anti-active Caspase 3 antibody (1:1000; Affinity purified Rabbit polyclonal, AF835; R&D) diluted in 1% BSA in TBST (TBS+0.5% triton). Next day, the sections were washed in TBS and incubated with secondary antibody (biotin labeled, 1:100 diluted with 1% BSA in TBST; Vector PK6200) for 90 minutes, followed by washing with TBS and incubation with streptavidin-biotin HRP labeled tertiary antibody for 90 minutes (SABC, 1:200 diluted in 1% TBST; Vector PK6200). The sections were further washed with TBS and treated with DAB solution for 20 minutes (0.025% 3, 3’-diaminobenzidine tetrahydrochloride; Sigma + 2.5% Nickel Sulphate Hexahydrate, Sigma+0.06% H_2_O_2_) for color development and visualization. The reaction was terminated under running tap water and the sections were rinsed with distilled water, dehydrated in alcohol series, cleared in xylene and mounted with distyrene plasticizer xylene (DPX) and stored for image analysis.

### Immunolabelling of β III tubulin and DCX proteins at PND 180

Hippocampal neurons expressing β III tubulin and DCX were detected using anti-β III tubulin and anti-DCX antibodies. Separate slides with hippocampal sections from each group (n=6 slides from different animals and dams/group) for both independent antibodies, were air dried and washed using PBS (Phosphate Buffer Saline). Washing was followed by permeabilization for 20 minutes using 0.5% Triton X-100 (Sigma) in PBS. The sections were then washed with PBST (0.1% Tween 20 in PBS), blocked in 1% H_2_O_2_ (Merck) and incubated with 1 % normal Serum (Vector PK6200) for 90 minutes. After non-specific blocking, the sections were incubated overnight at 4°C with primary antibodies (anti-β III tubulin mouse monoclonal, Sigma, T7816 and anti-DCX, Guinea Pig polyclonal, Millipore, AB5910) at dilution of 1:1000 in 1% BSA in PBST (0.5% triton added to PBS). Next day, the sections were washed in PBST for removal of unbound antibody and incubated with secondary antibody (biotin labeled, 1:100 diluted in 1% BSA in PBST; Vector PK6200) for 90 minutes, followed by washing with PBST and tertiary antibody incubation for 90 minutes (SABC, 1:200 dilution in 1% PBST; Vector PK6200). The sections were then washed with PBS and treated with DAB solution (0.025% 3, 3’-diaminobenzidine tetrahydrochloride; Sigma + 0.06% H_2_O_2_) for visualization. The reaction was terminated under running tap water, and sections were counterstained with hematoxylin (Vector) for proper visualization of the hippocampal layers. The sections were then dehydrated, cleared in xylene, mounted in DPX and stored for image analysis.

### Image analysis

For quantitative measurements, imageswith fixed frame size (21670.9 µm^2^) and magnification (20X) were grabbed separately for different regions of the hippocampus, using Leica DME 6000 microscope attached to a computer installed with LAS (Leica application suit) software. For quantitative analysis of different regions, data were analyzed separately, 18 images each for CA1, CA3 and DG were collected from a single slide and a total number of 6 slides from different animals were analyzed for every marker (n=108). During antigen density measurement, the percentage and intensity of positively labeled areas (area fraction) within the frame of all the images were individually detected using image quantification module of Leica Qwin software. The labeled area depicted the presence of positive cells as well the level of expression of the labeled protein by the respective cells. The mean percent area of immunostaining was calculated for CA1, CA3 and DG and was plotted as histograms. For cell count, images from different regions of hippocampus were quantified separately by interactive cell count module of Leica Qwin software and the number of cells were counted from each image/frame area and then calculated as number of cells/mm^2^ and plotted as mean cell count (Sharma et al., 2016).

### Golgi Technique

From each group, animals (n=6, belonging to different dams/group) were decapitated at PND 180, followed by dissection and immediate immersion of brain tissues in Golgi fixative solution (consists of sodium dichromate, chloral hydrate, formaldehyde, glutaraldehyde and DMSO). The tissues were immersed fixed for 72 hrs and then treated with 0.75% silver nitrate (Qualigens) solution for minimum 48 hrs. After complete impregnation, the sections (100µm thickness) were cut using a Leica Vibratome (VT 1000s). The sections were dehydrated, mounted in DPX and stored for neuronal analysis (Kumar et al., 2013).

### Morphological analysis of neurons

Neurons belonging to different animals from each group were traced using camera Lucida, attached to Leica DME microscope (n=36 neurons from 6 different slides/group). Sholl analysis for dendritic length and arborization was performed for each neuron, using ImageJ freeware software. The number of intersections in each Sholl circle was analyzed for complexity and length of neurites (Baharnoori et al., 2008).

### Cognitive experiment for memory analysis at PND 180 (n=12/group from different dams)

#### Morris Water Maze (MWM)

Spatial memory was assessed using MWM (Columbus Instruments) as per the procedure opted by Naik et al., (2015). Initially, the animals were acclimatized and taught to navigate the hidden/escape platform using spatial clues for three consecutive days (4 trial each animal for 120 second). 60 min of gap was given between the trials. During the learning period, if the animals failed to reach the platform, they were manually guided to the escape platform following the shortest and direct path from the start point to the escape platform. On the fourth day i.e., after 24 hrs, the animals were subjected to MWM test to locate the escape platform within 120 sec. Platform was removed on the fourth day to avoid visual error (probe test). Data was recorded for three trials using a vertical camera connected to an Any Maze software version 4.82. Parameters like the mean path efficiency i.e., the shortest route taken by the animal to reach the escape platform, latency i.e., the time taken to reach the escape platform and the percent time spent in the target quadrant (north-west) was analyzed for each and every group.

#### T Maze

For spatial working memory assessment, T maze from Columbus instrument was used and the protocol was designed according to Nagayach et al., (2014). It is a reward-based test, in which animals were trained and taught to detect the baited arm by remembering previous visited arm. It is a T shaped maze with one start and two reward arms. The animals were also made to learn left right discrimination in this test. Initially, the animals were acclimatized by placing in the maze and allowing them to explore the whole maze. During training, the animals were placed in the start arm and allowed to arbitrarily choose an arm for 30s. The animals were then removed and again placed on the start arm, expecting to choose the unexplored arm. The reward was placed at the end of either left or right arm and the position of the reward was changed after every trial. The animals were taught to obtain the reward making minimum effort while choosing the correct arm. 4 trials each of 120s were given to each animal with a gap of 60 min between trials for 3 consecutive days. On the 4^th^ day, 3 trials for each animal were performed and the mean path efficiency to reach the reward/baited arm was recorded using a vertical camera attached to an Any Maze software version 4.82.

#### Statistical Analysis

Data were analyzed using One Way ANOVA (for group wise comparison) and Two-Way ANOVA (for comparison between groups belonging to two independent variables i.e., diet and treatment) followed by post hoc Holm-Sidak test using Sigma Plot 12. Significance level was preset at P≤0.05.

The animals were maintained and experimental plan was designed with prior permission from Institutional Animal Ethics Committee of Jiwaji University, Gwalior (M.P), India.

## Abbreviations

HP: high protein;
LP: low protein;
PND: postnatal days;
Poly I:C: polyinosinic:polycytidylic acid;
LPS: Lipopolysaccharide;
HP+PolyI:C: highprotein+polyinosinic:polycytidylic acid;
HP+LPS: high protein+lipopolysaccharide;
HP+Poly I:C+LPS: high protein+polyinosinic:polycytidylic acid+lipopolysaccharide;
LP+Poly I:C: low protein+polyinosinic:polycytidylic acid;
LP+LPS: low protein+lipopolysaccharide;
LP+Poly I:C+LPS: low protein+polyinosinic:polycytidylic acid+lipopolysaccharide;
DCX: doublecortin;
Caspase: cysteine aspartic proteases;
MWM: Morris water maze;
CA: cornuammonis;
DG: dentate gyrus

## Acknowledgement

The authors are thankful to Indian Council of Medical Research (ICMR), New Delhi, India.

## Author contribution

Ishan Patro conceptualized and designed the study. Tiyasha Sarkar and Nisha Patro performed the experiments and data analysis and wrote the manuscript. Ishan Patro and Nisha Patro then finalized the manuscript.

## Compliance with Ethical Standards

All experiments on rats were performed in accordance with the Institutional Animal Ethics Committee of Jiwaji University and in compliance with National Institutes of Health Guide for the care and use of laboratory animals.

## Supplementary material

**Table 1**Statistical data of Sholl analysis

